# AICM: A Genuine Framework for Correcting Inconsistency Between Large Pharmacogenomics Datasets

**DOI:** 10.1101/386896

**Authors:** Zhiyue Tom Hu, Yuting Ye, Patrick A. Newbury, Haiyan Huang, Bin Chen

## Abstract

The inconsistency of open pharmacogenomics datasets produced by different studies limits the usage of pharmacogenomics in biomarker discovery. Investigation of multiple pharmacogenomics datasets confirmed that the pairwise sensitivity data correlation between drugs, or rows, across different studies (drug-wise) is relatively low, while the pairwise sensitivity data correlation between cell-lines, or columns, across different studies (cell-wise) is considerably strong. This common interesting observation across multiple pharmacogenomics datasets suggests the existence of subtle consistency among the different studies (i.e., strong cell-wise correlation). However, significant noises are also shown (i.e., weak drug-wise correlation) and have prevented researchers from comfortably using the data directly. Motivated by this observation, we propose a novel framework for addressing the inconsistency
between large-scale pharmacogenomics data sets. Our method can significantly boost the drug-wise correlation and can be easily applied to re-summarized and normalized datasets proposed by others. We also investigate our algorithm based on many different criteria to demonstrate that the corrected datasets are not only consistent, but also biologically meaningful. Eventually, we propose to extend our main algorithm into a framework, so that in the future when more data-sets become publicly available, our framework can hopefully offer a “ground-truth” guidance for references.

## 1. Introduction

One goal of precision medicine is to select optimal therapies for individual cancer patients based on individual molecular biomarkers identified from clinical trials.^1–3^ Molecular biomarkers for many cancer drugs are currently quite limited, and it takes many years to identify and validate a biomarker for a single drug in clinical trials.^4,5^ Recent pharmacogenomics studies, where drugs are tested against panels of molecularly characterized cancer cell lines, enabled large-scale identification of various types of molecular biomarkers by correlating drug sensitivity with molecular profiles of pre-treatment cancer cell lines.^6–10^ These biomarkers are expected to predict the chance that cancer cells will respond to individual drugs.

There have been a handful of similar pharmacogenomic studies since Cancer Cell Line Encyclopedia (CCLE)^7^ and Genomics of Cancer Genome Project (CGP)^11^ were published in2012 by the Broad Institute and Sanger Institute, respectively. CCLE included sensitivity data for 1046 cell lines and 24 compounds; CGP included data for almost 700 cell lines and 138 compounds. The following Broad Institute’s Cancer Therapeutics Response Portal (CTRPv2) dataset included 860 cell lines and 481 compounds.^8,12,13^ The dataset from the Institute for Molecular Medicine Finland (FIMM) included 50 cell lines and 52 compounds.^14^ The new version of Genomics of Drug Sensitivity in Cancer (GDSC1000) dataset included 1001 cell lines and 251 compounds. There have also been similar pharmacogenomics studies specific to particular cancers including acute myeloid leukemia.^15–17^

Each dataset is essentially a data matrix, where each row represents one drug, each column represent one cell-line, and values are sensitivity measures derived from dose–response curves. IC50 (concentration at which the drug inhibited 50% of the maximum cellular growth) and AUC (area under the activity curve measuring dose response) are commonly used as sensitivity measures. However, recent re-investigation of published pharmacogenomics data has revealed the inconsistency of drug sensitivity data among different studies, raising the concern of using them for biomarker discovery.^18,19^ In the recent comparison of drug sensitivity measures between CGP and CCLE for 15 drugs tested on the 471 shared cell lines, the vast majority of drugs yielded poor concordance (median Spearman’s rank correlation of 0.28 and 0.35 for IC50 and AUC, respectively).^18^

There have been numerous attempts to address this issue. Mpindi et al. proposed to increase the consistency through harmonizing the readout and drug concentration range.^20^ They re-analyzed the dose-response data using a standardized AUC response metric. They found high concordance between FIMM and CCLE and reasoned that similar experimental protocols were applied, including the same readout, similar controls. Bouhaddou et al. calculated a common viability metric across a shared log10-dose range, and computed slope, AUC values and found the new matrix could lead to better consistency.^21^ Hafner et al. proposed another metric called GR50 to summarize drug sensitivity and demonstrated its superiority in assessing the effects of drugs in dividing cells.^22^ Most proposed ideas focused on forming better summarization metric and/or standardizing experiments and data processing pipeline. Unfortunately, standardization methods cannot address the inconsistency issues of existing datasets, and re-summarization methods rely heavily on the assumption that the raw data is correct — while there surely exist some technical noises, as datasets produced under similar experimental protocols seem to be more consistent with each other.^20^ Hence when the overlapping part between datasets grows bigger and the noise sources become more complex, these methods might not work well. Note that most of the studies have focused on the overlaps between CCLE and other datasets, which only contain very limited number of drugs. Novel computational methods correcting large-scale summarized data are therefore in urgent need.

Studies confirmed that drug-wise correlation is poor, but the cell-wise correlation is considerably strong (for example: overlapping cell-lines between CTRPv2 and GDSC1000 have a median Spearman’s correlation of 0.553), suggesting the underlying consistency of pharmacogenomics datasets. Inspired by this observation, we developed a novel computational method Alternating Imputation and Correction Method (AICM). Through purely correcting data based on their cell-wise correlation, AICM significantly improves the drug-wise correlation and hence makes the datasets more credible in future work. We release the code and corrected datasets to the community^a^. To the best of our knowledge, this is the first method that leverages cell-wise information into correcting data to address such challenge.

## 2. Method

### 2.1. Method overview

The main goal is to increase the drug-wise correlation between two datasets, denoted as *A,B* ∈ ℝ^*n*×*p*^ — *n* drugs and *p* cell-lines — for convenience. We denote the *i*th **row** of matrix *A* as *A*_[*i*,:]_, then the goal can be formalized into the following problem:

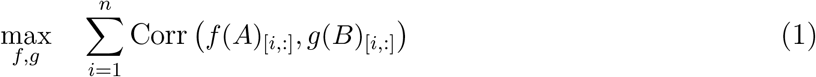

This is a more generalized idea than Renyi’s correlation as we define *f, g* not functions but **operations** such that *f, g* : ℝ^*n*×*p*^ → ℝ^*n*′×*p*′^, where *n′, p′* ∈ ℤ_+_. Operations include using a new summarization metric to re-summarize raw data and subsampling the data.

Now, since cell-wise correlation is consistently to be much more concordant across different studies than drug-wise correlation, we can raise one question: can we rely on the cell-wise information to correct the datasets so that the drug-wise correlation will also be improved? We denote *A*^*j*^ as *j*th **column** of *A* and *A^J^* as the union of all column *A^j^* such that *j* ∈ *J*, then more precisely, we want to develop some operation *f, g* such that

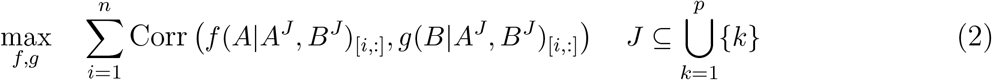

where 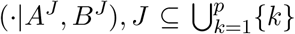 means either partial or all corresponding column information of *A* and *B* is given. We have found that there are considerably large amount of missing data in these datasets. Surprisingly, with some imputation of these missing data based solely on the cell-wise information, we found increase in drug-wise correlation. This confirmed our hypothesis that cell-wise information can be utilized to correct the datasets. Thus, AICM is developed to accomplish this goal by randomly dropping the parts of one dataset’s column and re-fit based on another dataset’s corresponding column with a simple linear regression with ℓ_*∞*_ norm regularization. The corrected values are subject to a hard threshold assuming that the data are not completely detained by noises, so that the corrected data shall not depart too far from the original value. By repeating such regression process interactively between two datasets, AICM hopes to reveal the true information shared in between these datasets and hence increase the drug-wise consistency.

### 2.2. Algorithm

The main idea is as described above: we uniformly randomly drop the values from one matrix (response matrix) and use the other matrix’s column (variable matrix) to impute dropped values. We then threshold the imputed values into the final correction by some proportional threshold with respect to the original values of the response matrix. We iteratively repeat this process by swapping the role of response and variable between two matrices. Below are the hyperparameters for the algorithm:

- max iterations (*iter* ∈ ℤ_+_): how many iterations the alternating imputation and correction need to be run.
- dropping rate (*r* ∈ (0,1)): what percent of the data from the response matrix should be dropped each iteration
- regularization term (*λ*_*r*_ ∈ ℝ_+_): how much the original value should be taken into account during the regression process
- hard proportional constraint (*λ*_*h*_ ∈ (0,1)): how many percentage points percent the imputed data can depart from the original value absolutely

And the full algorithm is described in detail as in Algorithm 1. Note that we use a simple linear regression with ℓ_*∞*_ norm (Eq 3) regularization for fitting process. Nevertheless, one can always use other fitting methods. For example, if one believes sparsity needs to be incorporated, one can use more weights and an ℓ_1_ norm, or if one believes there needs to be some group effects across cell-lines, one can use an ℓ_1_ and ℓ_2_ norm penalty. These ideas are similar to the idea of Lasso and Elastic Net.^23,24^ However, it is suggested that the objective function should remain convex, since solving non-convex problems would highly likely lead to a local extrema (or even a saddle point) and thus cause disastrous variations among trials.

### 2.3. Remarks

The objective function (3) is convex and hence would always achieve a global minimum with an appropriate solver. The problem can be solved efficiently and accurately by various methods such as proximal gradient algorithm and alternating direction of multipliers (ADMM).^25,26^ They have well-established convergence theorems and are available in many open-source (i.e. SCS^27^) and industrial solvers.^28^

In the next section, we will show the results of our algorithm on real datasets, as well as synthetic datasets to demonstrate our method significantly increases drug-wise correlation remarkably and does not artificially increase the correlation under certain assumption. We will also show the result is indeed biologically meaningful.

## 3. Results and Discussion

### 3.1. Synthetic datasets

The alternative correction procedure (**Swap**) in AICM essentially agglomerates two datasets. It inevitably gives rise to the concern that the corrected datasets are forced to be similar regardless of the ground truth. For example, one easily questions whether AICM improves the between-group correlation of placebo – it functions as white noise, thus is expected to be uncorrelated between one dataset and another. In addition, the induced randomness (Drop) in AICM might well shake one’s confidence in the stability and reliability of this method. In this section, we utilize synthetic datasets to demonstrate that AICM are free of these hypothetical troubles.

#### Algorithm 1 Alternating Imputation and Correction Method (AICM)

**Hyperparameter**: Dropping rate *r*, maximum iteration *iter*, regularization term *λ*_*r*_, and hard constraint term *λ*_*h*_.

**Input**: Two data matrices, of both *n* drugs and *p* cell-lines with summarized sensitivity data, denote as *A, B* ∈ ℝ^*n*×*p*^. We denote *j*th column of two matrices as *a*^*j*^, *b*^*j*^, *j* ∈ {1, 2,…, *p*} respectively. We denote the entry at *i*th row and *j*th column as *A_ij_* and *B_j_* respectively, {*i*,*j*} ∈ {1,2,…, *n*} × {1,2,…, *p*}.

**Initialization**: For each *j* ∈ {1, 2,…, *p*}, for all *i* ∈ {1,2,… *n*} such that *B_ij_* is missing while *A_ij_* is not, we denote such set as 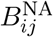, we fit a linear model such that *α*_*j*_, *β*_*j*_ maximizes ‖*b^j^* = *α_j_a^j^* + *β_j_*‖_2_ and then impute the missing values as 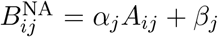. Then swap the role of *A* and *B* and repeat the above process. Now we have two matrices with same missing indices.

**for** *k* in {1,2,…*Iter*} **do**

**Swap**: *A* → *B*, *B* → *A*.

**Drop**: Randomly drop *r* × *n* × *p* data uniformly from *A*, we denote the indices of the dropped data as 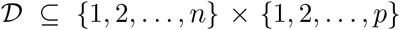, and hence dropped data as a set 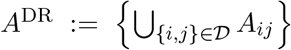. In a similar fashion, we denote dropped data of **column** *k* as 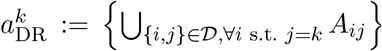, we denote the corresponding data in *k*th column of *B* as 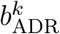. We fit a set of parameters *α_j_* ∈ ℝ, *β_j_* ∈ ℝ for each *j* with the following objective function:

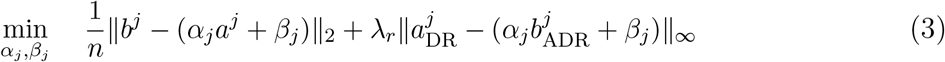

**Correction**: Set 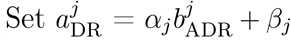 for each *j*. We denote the set of corrected value as 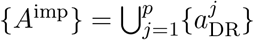.

**Threshold**: For 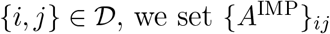 to

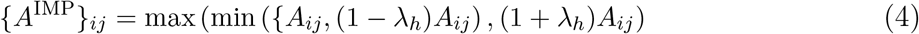

**end for**

In the most ideal scenario, where there exist no technical or biological noises, the drug sensitivity matrices are expected to be the same across distinct research teams. For simplicity, we assume that the ground truth can be separated into the drug part and the cell part. Then, the observed matrix can be modelled as

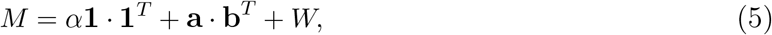

where *α* is the baseline, **a** ∈ ℝ^*n*^ contains the information about the *n* drugs, *b* ∈ ℝ^p^ summarizes the structure of the cell lines. The matrix α**1** · **1**^*T*^ + **a · b**^*T*^ represents the ground truth of the drug sensitivities. We simulate the ineffective drugs as uncorrelated rows by setting the top m entries of a to 0’s while the other rows associated with non-zero values (hence correlated) in a are regarded as effective drugs. *W* ∈ ℝ^*n*^×^*p*^ is a random matrix from a matrix normal distribution which reflects the composite of noise. In this study, we set *n* = 50, *p* = 40, *m* = 10.

The details of the data generation process are deferred to supplementary material^b^.

We apply AICM to the synthetic datasets with 30 different combinations of hyperparameters iter and *λ*_*h*_: *iter* ∈ {20,40,80,100,120,140} and *λ*_*h*_ ∈ {0.05,0.1,0.15,0.2,0.25}, and repeat the method for 20 times for each combination. With careful selection, we take (*iter*, *λ*_*h*_) = (80,0.1) because this combination gives acceptable reduction on correlations between first ten uncorrelated rows and strong increase of correlations between correlated rows as demonstrated (see Figure 1). In addition, *λ*_*h*_ = 0.1 is a conservative control of the correction step. Note that the normalized distances between the two matrices and the ground truth are reduced to 1.188 and 1.170 respectively after correction (the distances are 1.272 and 1.267 before correction). The decrease in distance is relatively significant, given the fact that we put a hard proportional threshold at 10% for each individual value. Therefore, AICM does help reduce the noise in the observed matrices. Furthermore, the Spearman’s correlation median of the correlated rows is increased to 0.390 from 0.219 with standard deviation 0.021, while the Spearman’s correlation median of uncorrelated rows is reduced to 0.084 from 0.095 with standard deviation 0.010. It indicates that the result is insensitive to the randomness of the dropping procedure in AICM. In Figure 2, the actual shift of the correlation distributions is displayed. On top of incremental correlations of correlated rows, there appear to be reduced correlations of uncorrelated rows after using AICM. It implies that our method not only enhances the real signals, but also exposes the fake ones. Thus, the original concern is eliminated on indiscriminately blending signals between datasets.

**Fig. 1:**
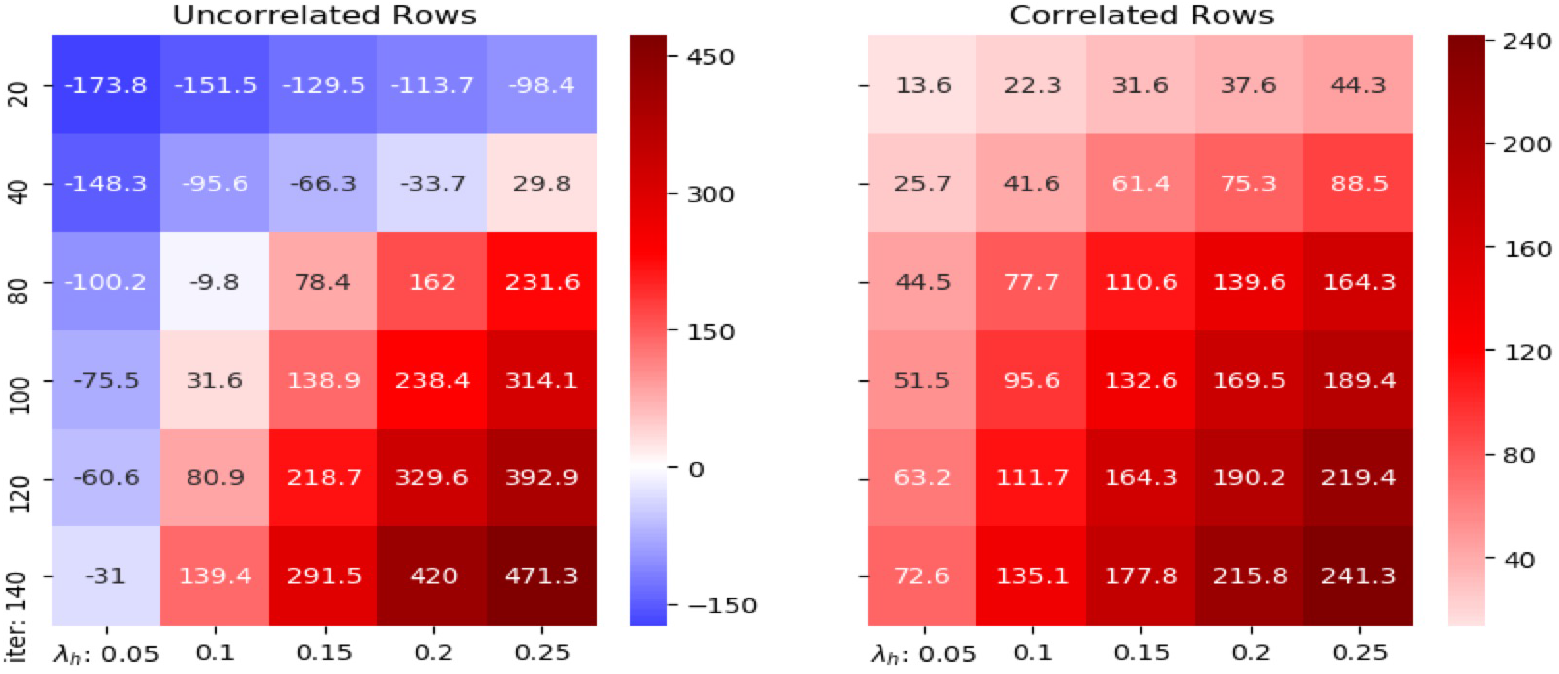
The percentage change (%) of the medians of the correlations on synthetic datasets with different parameters. x-axis is iter and y-axis is λ_h_.

**Fig. 2:**
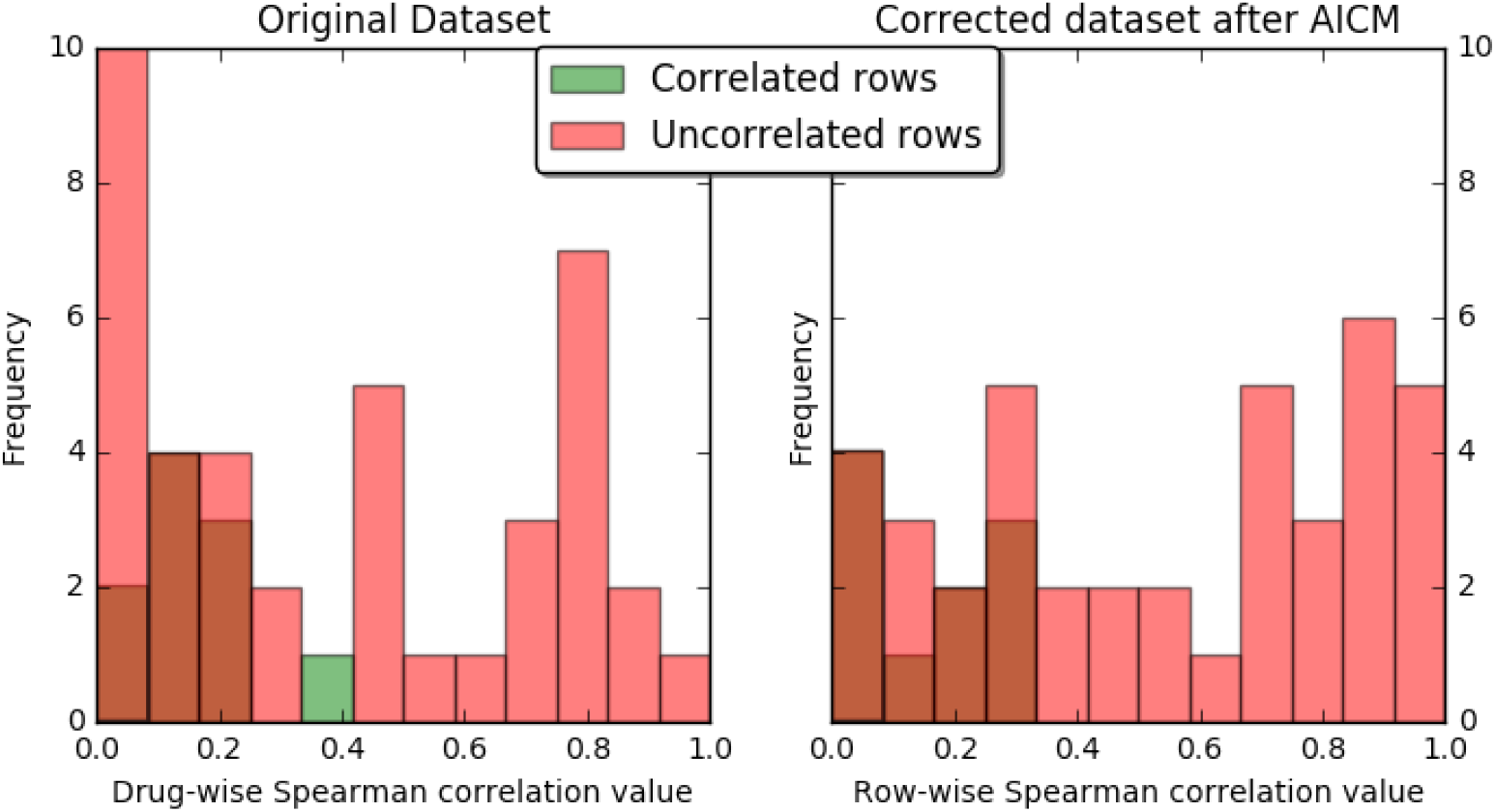
Distribution of drug-wise correlations between the synthetic datasets before AICM is applied and after.

### 3.2. Real datasets

We chose the three largest datasets in PharmacoGX: CTRPv2, GDSC1000, and FIMM as case studies.^8,11,13,19^ Drug names were compared by first converting to InChIKey via the webchem

R package.^29^ For the GDSC1000 dataset, 60 InChIKeys were subsequently manually retrieved from PubChem. A Python script was prepared and used to retrieve generic cell line “Accession numbers” from Cellosaurus.^30^ Given that not all cell lines returned Accession numbers, we removed symbols, spaces, and case from the names of the remaining cell lines for improved matching between datasets. For each of the three datasets, their respective IC50 and AUC data were obtained from PharmacoGx. Duplicate experiments were removed from CTRPv2 and GDSC1000 by removing all instances of a certain culture medium. Finally, the six dataframes were filtered for matching cell lines and drugs between each other, yielding 12 dataframes which contain IC50 and AUC between all 3 datasets.

With the optimal hyperparameters fetched from synthetic data, we demonstrate the shift of Spearman’s correlation between 90 drugs overlapping between GDSC1000 and CTRPv2 after AICM is deployed in Figure 3a. The data uses AUC summarization. It is clear that after AICM is deployed, the two datasets become more concordant with each other — this can be observed from both individual drug scatter plot and overall distribution. We also demonstrate two similar graphs between 30 overlapping drugs between CTRPv2 and FIMM, 29 overlapping drugs between GDSC1000 and FIMM with AUC summarization in Figure 3b and 3c.

**Fig. 3:**
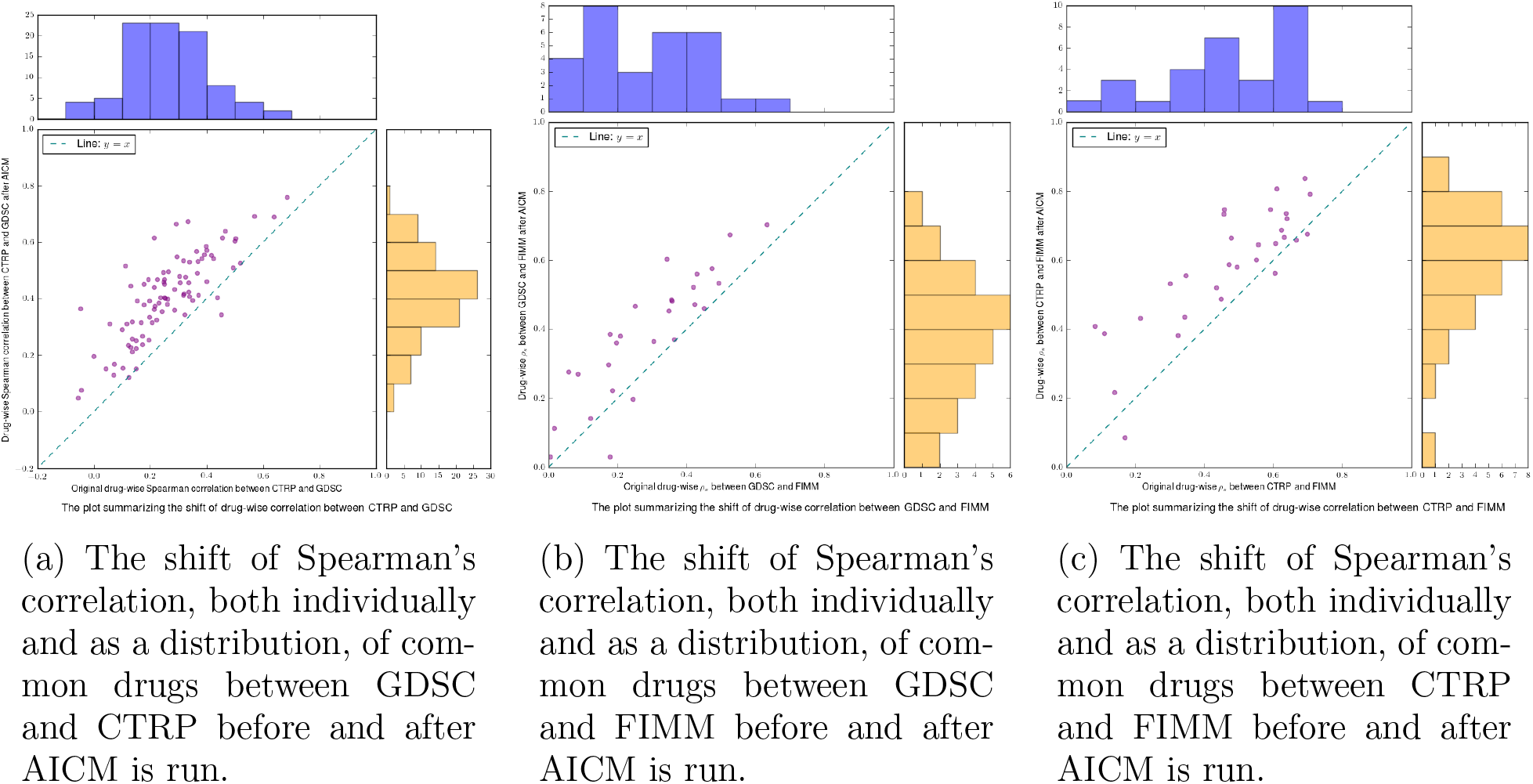
Summarization of individual drugs’ improvement and overall marginal distribution shift between FIMM, CTRP and GDSC.

Note that when we calculate the correlation, the original values that are missing are discarded from both matrices for fair comparison. Brief statistics of the original and post-correction drug-wise Spearman’s correlation can be found in Table 1. For significance, we used the cutoff of one-sided test at *p*-value 0.05 using the significance test of Spearman’s correlation proposed by Jerrold Zar.^31^ The values present what percentage of drugs is significant across two datasets.

**Table 1:** Brief statistics of the original and post-correction drug-wise Spearman’s correlation.

| Datasets | Mean |  | Median |  | Significant |  | Size |  |
| --- | --- | --- | --- | --- | --- | --- | --- | --- |
|  | Before | After | Before | After | Before | After | Drug | Cell |
| CTRPv2 & GDSC1000 | 0.261 | 0.410 | 0.249 | 0.411 | 63.33% | 90.00% | 90 | 566 |
| CTRPv2 & FIMM | 0.485 | 0.624 | 0.468 | 0.585 | 70.00% | 93.33% | 30 | 41 |
| GDSC1000 & FIMM | 0.250 | 0.352 | 0.278 | 0.380 | 27.59% | 55.17% | 29 | 47 |

We also demonstrate the scatter plots of some individual drug’s effect on cell-lines before and after AICM correction in Figure 4, we can indeed see the scatter plots become more concordant across datasets. We color the plots in a similar fashion as Safikhani et al.: we use blue (sensitive) to denote both datasets ≥ 0.2 and red (resistant) for both ≤ 0.2; orange denotes inconsistency.^19^ We pay particular interest to drugs that show significant improvement and drugs that show little improvement. We can see that drugs such as ZSTK474. Rapamycin, JQ1, OSI027 and PIK93 show significant improvement. Although Velaparib shows little improvement, it is known to be a very selective PARP inhibitor; it is not effective in any of cancer cell-lines examined in this study. Thus it would be meaningless and artificial to increase the correlation across two datasets.

We also present the scatter plots of some drugs shared by all three datasets: CTRPv2, GDSC1000 and FIMM. We can see that in both 5a and 5b, the two graphs on the right consistently demonstrate more similar pattern than the two graphs on the left, which confirms that the variation across multiple datasets is alleviated after AICM is deployed – AICM indeed recovers some meaningful signals.

**Fig. 4:**
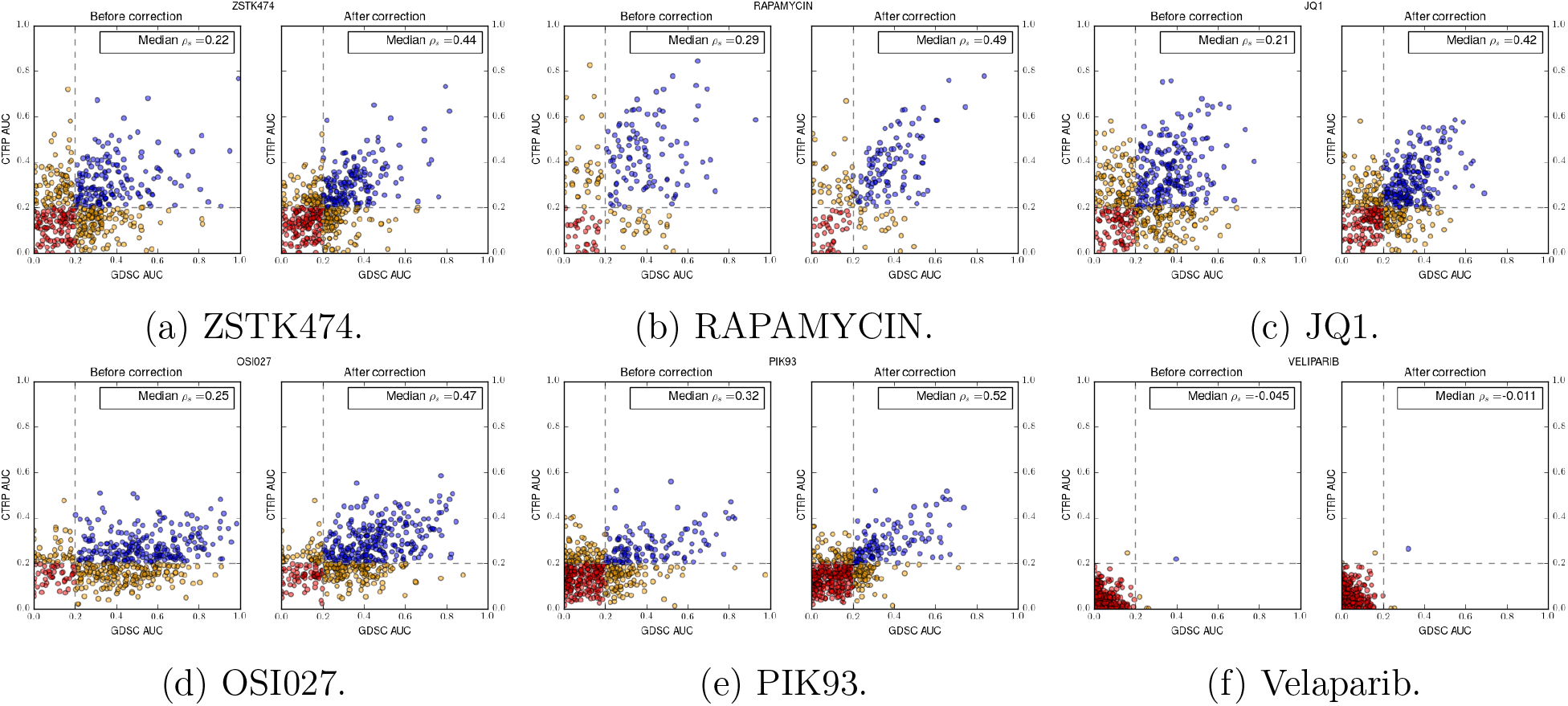
Individual drugs with respect to individual cell-lines before and after AICM is deployed. First five demonstrate drugs whose correlations are significantly improved and the last one demonstrates a drug whose correlation is poorly improved.

**Fig. 5:**
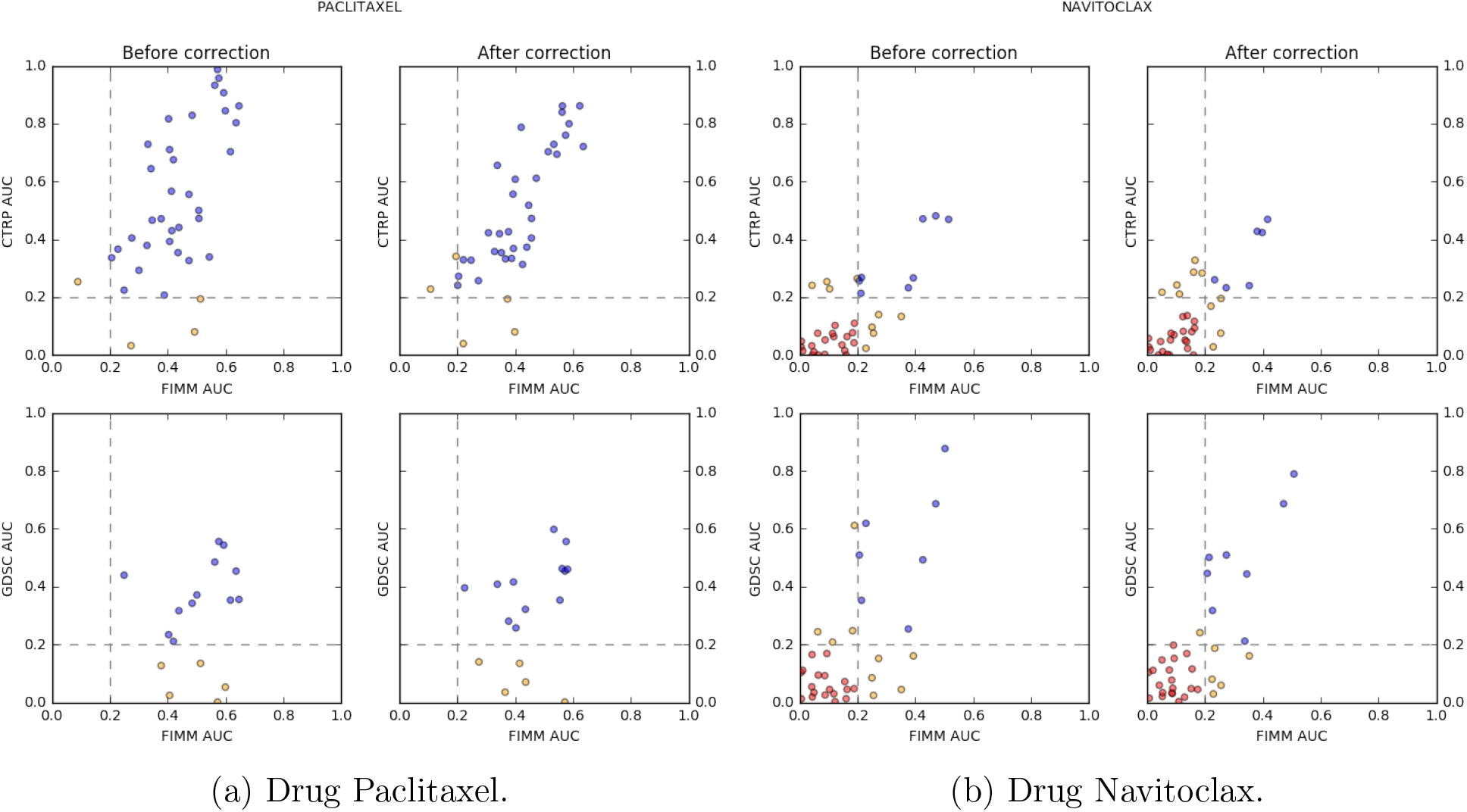
Overlapping drugs across three datasets.

## 4. Conclusions and Future Work

In this work, we develop a genuine algorithm by alternatively dropping and fitting cell-wise data and succeds in improving the drug-wise correlation. The algorithm can easily incorporate other different ideas. For example, one can replace the fitting process with other regression methods if one had prior assumptions in mind. We have shown that with appropriate hyperparameters chosen, AICM can improve the drug-wise correlation across different studies and that the increase in correlation is indeed concordant and biologically meaningful.

We realize the limitation of AICM’s dependence on the overlapping of existing data, while such data is rather rare. We did not include experiment on CCLE dataset primarily because it has very limited drug overlap with other existing datasets. Also, AICM currently does not purport to correct sensitivity data of new drugs. Future work will be to extend such algorithm into a complete framework. We will maintain a database of corrected existing drugs and cells, and when more data come in, we will be able to incorporate it. We hope as more data comes in, the database would asymptotically become more accurate of reflecting true relationship between drugs and cell-lines and can thus serve as a ground-truth guidance. As for new drugs, we will develop either a generative algorithm or a clustering algorithm, i.e. getting the latent distribution where drug is “generated” or cluster it based on existing features, and find similar existing drugs in hope of some practical guidance. We believe our corrected datasets will facilitate biomarker discovery.

## Acknowledgments

Preprint of an article submitted for consideration in Pacific Symposium on Biocomputing © 2018, copyright World Scientific Publishing Company, http://psb.stanford.edu. This work is supported by R21 TR001743 and K01 ES028047 and the MSU Global Impact Initiative. We would thank Anthony Sciarini for providing the pipeline to fetch the cell-line generic names. We would also thank Ryan Lovett and Chris Paciorek for all helps received on cluster computing issues.

## Footnotes

a www.github.com/tomwhoooo/aicm

b www.github.com/tomwhoooo/aicm/paper_supp

## References

1. F. S. Collins and H. Varmus. A new initiative on precision medicine. N. Engl. J. Med., 372(9):793–795, Feb 2015.

2. D. R. Lowy and F. S. Collins. Aiming High–Changing the Trajectory for Cancer. N. Engl. J. Med., 374(20):1901–1904, May 2016.

3 B. Chen and A. J. Butte. Leveraging big data to transform target selection and drug discovery. Clin. Pharmacol. Ther., 99(3):285–297, Mar 2016.

4. G. Yothers, M. J. O’Connell, M. Lee, M. Lopatin, K. M. Clark-Langone, C. Millward, S. Paik, S. Sharif, S. Shak, and N. Wolmark. Validation of the 12-gene colon cancer recurrence score in NSABP C-07 as a predictor of recurrence in patients with stage II and III colon cancer treated with fluorouracil and leucovorin (FU/LV) and FU/LV plus oxaliplatin. J. Clin. Oncol., 31(36):4512–4519, Dec 2013.

5. A. de Gramont, S. Watson, L. M. Ellis, J. Rodon, J. Tabernero, A. de Gramont, and S. R. Hamilton. Pragmatic issues in biomarker evaluation for targeted therapies in cancer. Nat Rev Clin Oncol, 12(xyi4):197–212, Apr 2015.

6. M. J. Garnett, E. J. Edelman, S. J. Heidorn, C. D. Greenman, A. Dastur, K. W. Lau, P. Greninger,L.R. Thompson, X. Luo, J. Soares, Q. Liu, F. Iorio, D. Surdez, L. Chen, R. J. Milano, G. R. Bignell, A. T. Tam, H. Davies, J. A. Stevenson, S. Barthorpe, S. R. Lutz, F. Kogera, K. Lawrence, A. McLaren-Douglas, X. Mitropoulos, T. Mironenko, H. Thi, L. Richardson, W. Zhou, F. Jewitt,T. Zhang, P. O’Brien, J. L. Boisvert, S. Price, W. Hur, W. Yang, X. Deng, A. Butler, H. G. Choi, J. W. Chang, J. Baselga, I. Stamenkovic, J. A. Engelman, S. V. Sharma, O. Delattre, J. Saez-Rodriguez, N. S. Gray, J. Settleman, P. A. Futreal, D. A. Haber, M. R. Stratton, S. Ramaswamy, U. McDermott, and C. H. Benes. Systematic identification of genomic markers of drug sensitivity in cancer cells. Nature, 483(7391):570–575, Mar 2012.

7. J. Barretina, G. Caponigro, N. Stransky, K. Venkatesan, A. A. Margolin, S. Kim, C. J. Wilson, J. Lehar, G. V. Kryukov, D. Sonkin, A. Reddy, M. Liu, L. Murray, M. F. Berger, J. E. Monahan, P. Morais, J. Meltzer, A. Korejwa, J. Jane-Valbuena, F. A. Mapa, J. Thibault, E. Bric-Furlong, P. Raman, A. Shipway, I. H. Engels, J. Cheng, G. K. Yu, J. Yu, P. Aspesi, M. de Silva, K. Jagtap, M. D. Jones, L. Wang, C. Hatton, E. Palescandolo, S. Gupta, S. Mahan, C. Sougnez, R. C. Onofrio, T. Liefeld, L. MacConaill, W. Winckler, M. Reich, N. Li, J. P. Mesirov, S. B. Gabriel, G. Getz, K. Ardlie, V. Chan, V. E. Myer, B. L. Weber, J. Porter, M. Warmuth, P. Finan, J. L. Harris, M. Meyerson, T. R. Golub, M. P. Morrissey, W. R. Sellers, R. Schlegel, and L. A. Garraway. The Cancer Cell Line Encyclopedia enables predictive modelling of anticancer drug sensitivity. Nature, 483(7391):603–607, Mar 2012.

8. A. Basu, N. E. Bodycombe, J. H. Cheah, E. V. Price, K. Liu, G. I. Schaefer, R. Y. Ebright, M. L. Stewart, D. Ito, S. Wang, A. L. Bracha, T. Liefeld, M. Wawer, J. C. Gilbert, A. J. Wilson, N. Stransky, G. V. Kryukov, V. Dancik, J. Barretina, L. A. Garraway, C. S. Hon, B. Munoz, J. A. Bittker, B. R. Stockwell, D. Khabele, A. M. Stern, P. A. Clemons, A. F. Shamji, and S. L. Schreiber. An interactive resource to identify cancer genetic and lineage dependencies targeted by small molecules. Cell, 154(5):1151–1161, Aug 2013.

9. F. Iorio, T. A. Knijnenburg, D. J. Vis, G. R. Bignell, M. P. Menden, M. Schubert, N. Aben, E. Goncalves, S. Barthorpe, H. Lightfoot, T. Cokelaer, P. Greninger, E. van Dyk, H. Chang, G. de Silva, H. Heyn, X. Deng, R. K. Egan, Q. Liu, T. Mironenko, X. Mitropoulos, L. Richardson, J. Wang, T. Zhang, S. Moran, S. Sayols, M. Soleimani, D. Tamborero, N. Lopez-Bigas, P. Ross-Macdonald, M. Esteller, N. S. Gray, D. A. Haber, M. R. Stratton, C. H. Benes, L. F.A. Wessels, J. Saez-Rodriguez, U. McDermott, and M. J. Garnett. A Landscape of Pharmacogenomic Interactions in Cancer. Cell, 166(3):740–754, Jul 2016.

10. M. Niepel, M. Hafner, E. A. Pace, M. Chung, D. H. Chai, L. Zhou, B. Schoeberl, and P. K. Sorger. Profiles of Basal and stimulated receptor signaling networks predict drug response in breast cancer lines. Sci Signal, 6(294):ra84, Sep 2013.

11. W. Yang, J. Soares, P. Greninger, E. J. Edelman, H. Lightfoot, S. Forbes, N. Bindal, D. Beare, J. A. Smith, I. R. Thompson, S. Ramaswamy, P. A. Futreal, D. A. Haber, M. R. Stratton, C. Benes, U. McDermott, and M. J. Garnett. Genomics of Drug Sensitivity in Cancer (GDSC): a resource for therapeutic biomarker discovery in cancer cells. Nucleic Acids Res., 41(Database issue):D955–961, Jan 2013.

12. M. G. Rees, B. Seashore-Ludlow, J. H. Cheah, D. J. Adams, E. V. Price, S. Gill, S. Javaid, M. E. Coletti, V. L. Jones, N. E. Bodycombe, C. K. Soule, B. Alexander, A. Li, P. Montgomery, J. D. Kotz, C. S. Hon, B. Munoz, T. Liefeld, V. Dan?ik, D. A. Haber, C. B. Clish, J. A. Bittker, M. Palmer, B. K. Wagner, P. A. Clemons, A. F. Shamji, and S. L. Schreiber. Correlating chemical sensitivity and basal gene expression reveals mechanism of action. Nat. Chem. Biol., 12(2):109–116, Feb 2016.

13. B. Seashore-Ludlow, M. G. Rees, J. H. Cheah, M. Cokol, E. V. Price, M. E. Coletti, V. Jones, N. E. Bodycombe, C. K. Soule, J. Gould, B. Alexander, A. Li, P. Montgomery, M. J. Wawer, N. Kuru, J. D. Kotz, C. S. Hon, B. Munoz, T. Liefeld, V. Dan?ik, J. A. Bittker, M. Palmer, J. E. Bradner, A. F. Shamji, P. A. Clemons, and S. L. Schreiber. Harnessing Connectivity in a Large-Scale Small-Molecule Sensitivity Dataset. Cancer Discov, 5(11):1210–1223, Nov 2015.

14. B. Yadav, T. Pemovska, A. Szwajda, E. Kulesskiy, M. Kontro, R. Karjalainen, M. M. Majumder, D. Malani, A. Murumagi, J. Knowles, K. Porkka, C. Heckman, O. Kallioniemi, K. Wennerberg, and T. Aittokallio. Quantitative scoring of differential drug sensitivity for individually optimized anticancer therapies. Sci Rep, 4:5193, Jun 2014.

15. R. Marcotte, A. Sayad, K. R. Brown, F. Sanchez-Garcia, J. Reimand, M. Haider, C. Virtanen, J. E. Bradner, G. D. Bader, G. B. Mills, D. Pe’er, J. Moffat, and B. G. Neel. Functional Genomic Landscape of Human Breast Cancer Drivers, Vulnerabilities, and Resistance. Cell, 164(1-2):293–309, Jan 2016.

16. A. Daemen, O. L. Griffith, L. M. Heiser, N. J. Wang, O. M. Enache, Z. Sanborn, F. Pepin, S. Durinck, J. E. Korkola, M. Griffith, J. S. Hur, N. Huh, J. Chung, L. Cope, M. J. Fackler, C. Umbricht, S. Sukumar, P. Seth, V. P. Sukhatme, L. R. Jakkula, Y. Lu, G. B. Mills, R. J. Cho, D.A. Collisson, L. J. van’t Veer, P. T. Spellman, and J. W. Gray. Modeling precision treatment of breast cancer. Genome Biol., 14(10):R110, 2013.

17. S. I. Lee, S. Celik, B. A. Logsdon, S. M. Lundberg, T. J. Martins, V. G. Oehler, E. H. Estey, C. P. Miller, S. Chien, J. Dai, A. Saxena, C. A. Blau, and P. S. Becker. A machine learning approach to integrate big data for precision medicine in acute myeloid leukemia. Nat Commun, 9(1):42, 01 2018.

18. B. Haibe-Kains, N. El-Hachem, N. J. Birkbak, A. C. Jin, A. H. Beck, H. J. Aerts, and J. Quackenbush. Inconsistency in large pharmacogenomic studies. Nature, 504(7480):389–393, Dec 2013.

19. Z. Safikhani, P. Smirnov, M. Freeman, N. El-Hachem, A. She, Q. Rene, A. Goldenberg, N. J. Birkbak, C. Hatzis, L. Shi, A. H. Beck, H. J. W. L. Aerts, J. Quackenbush, and B. Haibe-Kains. Revisiting inconsistency in large pharmacogenomic studies. F1000Res, 5:2333, 2016.

20. John Patrick Mpindi, Bhagwan Yadav, Päivi Östling, Prson Gautam, Disha Malani, Astrid Murumägi, Akira Hirasawa, Sara Kangaspeska, Krister Wennerberg, Olli Kallioniemi, and Tero Aittokallio. Consistency in drug response profiling. Nature, 540:E5 EP-, 11 2016.

21. Mehdi Bouhaddou, Matthew S. DiStefano, Eric A. Riesel, Emilce Carrasco, Hadassa Y. Holzapfel, DeAnalisa C. Jones, Gregory R. Smith, Alan D. Stern, Sulaiman S. Somani, T. Victoria Thompson, and Marc R. Birtwistle. Drug response consistency in ccle and cgp. Nature, 540:E9 EP-, 11 2016.

22. M. Hafner, M. Niepel, M. Chung, and P. K. Sorger. Growth rate inhibition metrics correct for confounders in measuring sensitivity to cancer drugs. Nat. Methods, 13(6):521–527, 06 2016.

23. Hui Zou and Trevor Hastie. Regularization and variable selection via the elastic net. Journal of the Royal Statistical Society, Series B, 67:301–320, 2005.

24. Robert Tibshirani. Regression shrinkage and selection via the lasso. Journal of the Royal Statistical Society. Series B (Methodological), 58(1):267–288, 1996.

25. Neal Parikh and Stephen Boyd. Proximal algorithms. Found. Trends Optim., 1(3):127–239, January 2014.

26. Stephen Boyd, Neal Parikh, Eric Chu, Borja Peleato, and Jonathan Eckstein. Distributed optimization and statistical learning via the alternating direction method of multipliers. Found. Trends Mach. Learn., 3(1):1–122, January 2011.

27. B. O’Donoghue, E. Chu, N. Parikh, and S. Boyd. SCS: Splitting conic solver, version 2.0.2, November 2017.

28. Tianyi Lin, Shiqian Ma, and Shuzhong Zhang. On the global linear convergence of the admm with multiblock variables. SIAM Journal on Optimization, 25:1478–1497, 2015.

29. George Nicola, Tiqing Liu, and Michael K. Gilson. Public domain databases for medicinal chemistry. Journal of Medicinal Chemistry, 55(16):6987–7002, 2012. PMID: 22731701.

30. A. Bairoch. The Cellosaurus, a Cell-Line Knowledge Resource. J Biomol Tech, May 2018.

31. Jerrold H. Zar. Significance testing of the spearman rank correlation coefficient. Journal of the American Statistical Association, 67:578–580, 1972.

